# Engineered Heterochronic Parabiosis in 3D Microphysiological System for Identification of Muscle Rejuvenating Factors

**DOI:** 10.1101/2020.03.10.975482

**Authors:** Yunki Lee, Jeongmoon J. Choi, Song Ih Ahn, Nan Hee Leea, Woojin M. Han, Mahir Mohiuddin, Eun Jung Shin, Levi Wood, Ki Dong Park, YongTae Kim, Young C. Jang

**Affiliations:** School of Biological Sciences, Georgia Institute of Technology, Atlanta, GA 30332, USA; George W. Woodruff School of Mechanical Engineering, Georgia Institute of Technology, Atlanta, GA 30332, USA; Wallace H. Coulter Department of Biomedical Engineering, Georgia Institute of Technology, Atlanta, GA 30332, USA; Parker H. Petit Institute for Bioengineering and Bioscience, Georgia Institute of Technology, Atlanta, GA 30332, USA; Institute for Electronics and Nanotechnology, Georgia Institute of Technology, Atlanta, GA 30332, USA; Department of Molecular Science and Technology, Ajou University, Suwon 16499, Republic of Korea

**Keywords:** parabiosis, aging, muscle stem cell, muscle stem cell niche, microfluidics, organon-a-chip

## Abstract

Exposure of aged mice to a young systemic milieu revealed remarkable rejuvenation effects on aged tissues, including skeletal muscle. Although some candidate factors have been identified, the exact identity and the underlying mechanisms of putative rejuvenating factors remain elusive, mainly due to the complexity of *in vivo* parabiosis. Here, we present an *in vitro* muscle parabiosis system that integrates young- and old-muscle stem cell vascular niche on a three-dimensional microfluidic platform designed to recapitulate key features of native muscle stem cell microenvironment. This innovative system enables mechanistic studies of cellular dynamics and molecular interactions within the muscle stem cell niche, especially in response to conditional extrinsic stimuli of local and systemic factors. We demonstrate that vascular endothelial growth factor (VEGF) signaling from endothelial cells and myotubes synergistically contribute to the rejuvenation of the aged muscle stem cell function. Moreover, with the adjustable on-chip system, we can mimic both blood transfusion and parabiosis and detect the time-varying effects of anti-geronic and pro-geronic factors in a single organ or multi-organ systems. Our unique approach presents a complementary *in vitro* model to supplement *in vivo* parabiosis for identifying potential anti-geronic factors responsible for revitalizing aging organs.

## Introduction

Skeletal muscle possesses a remarkable ability to regenerate following injury due to the presence of resident muscle stem cell (MuSC), also known as satellite cell (1). MuSCs play an indispensable role in the maintenance of muscle homeostasis and repair through adult myogenesis (2, 3). Under physiological conditions, they remain dormant in quiescence underneath the basal lamina and adjacent to myofibers (1, 4). However, when stimulated by muscle damage or stress, the quiescent MuSCs undergo asymmetric division to either self-renew or activate a set of transcriptional factors for proliferation and myogenic differentiation and then fuse with existing myofibers to complete muscle regeneration process (5, 6). A growing amount of evidence suggests that this myogenic process is also influenced by a dynamic interplay between intrinsic factors within MuSCs and cellular/acellular extrinsic factors constituting the niche such as microvasculature and myofiber (4, 5, 7, 8). However, similar to other somatic stem cells, the regenerative capacity significantly declines with aging, leading to a slow and incomplete recovery process (7, 9-12). Although the MuSCs continue to enter the cell cycle and activate, their self-renewal capacity is compromised due to the aging-triggered stress responses such as DNA damage, oxidative stress, or cellular senescence, thus resulting in a reduction in function and quantity of quiescent MuSC pool (13-16). These events contribute to the loss of muscle integrity and a decline in muscle strength, commonly referred to as sarcopenia (11). Consequently, age-acquired deficits in muscle function influence quality of life in the elderly, and also directly correlate to morbidity and mortality (17).

In the past years, we and others have demonstrated that remarkable rejuvenating effects can be achieved in aged tissues including muscle, heart, brain, bone, and liver by surgically joining young and old mice through heterochronic parabiosis (15, 18-21). Intriguingly, exposure of aged mice to a young systemic microenvironment restored the regenerative capacity of endogenous MuSCs in the aged muscle, rejuvenating the aged muscle function and structure (15, 19, 22). Although the exact mechanisms remain to be understood, the continuous access to young blood and secretory organs is thought to boost immune function and attenuate inflammation by releasing humoral factors in the circulation (23, 24). Moreover, recent studies revealed potential systemic factors (*e.g.*, GDF11 and oxytocin) in young blood that are responsible for anti-geronic effects of parabiosis, demonstrating their rejuvenation effects on the aged muscle (25-27). Despite the tremendous therapeutic promise, the identity and underlying biochemical and cellular mechanisms of blood-borne factors regulating regeneration and homeostasis of skeletal muscle have yet to be elucidated. This gap in knowledge is in part due to the complexity of *in vivo* parabiosis and the dynamic nature of blood-borne factors (28, 29). Thus, designing a mechanistic *in vivo* study to verify the precise mechanisms of rejuvenation has been a major challenge.

Advances in microfluidics combined with cellular engineering have demonstrated the successful applications to serve as a complementary *in vitro* model of tissue-level human physiology called organ-on-a-chip (30, 31). Compared to conventional *in vivo* and *in vitro* experimental models, precise control of small-volume samples in organ-on-a-chip systems facilitates highly sensitive analysis to identify and validate critical soluble factors in *in vivo*-like organ-specific microenvironment (30, 32-34). Recent attempts to integrate organ-on-a-chip platforms have provided novel approaches to investigate complex cell-cell communications and dynamics between different physiological functional units (33, 35). Capitalizing on this integrated microphysiological system, we can simulate organ-specific or multi-organ parabiosis with modulated inter-circulation of humoral factors in the local cellular microenvironment in a closed-loop circulatory system. Furthermore, a high-precision control technology incorporated with microfluidic systems enables real-time monitoring and identification of secretory factors (36, 37).

Here, we report a muscle parabiosis-on-a-chip system that harnesses biochemical and biophysical properties of the MuSC vascular microenvironment and inter-circulation of blood-borne-factors between young and old MuSC microenvironments. We first engineer a 3D vascularized MuSC niche-on-a-chip (VMoC) device that closely reconstitutes the architecture of the native MuSC microenvironment. We then demonstrate the myogenic activities of MuSCs in response to conditional extrinsic stimuli (*e.g.*, local and systemic environments), and validate the crosstalk between MuSCs and ECs *via* vascular endothelial growth factor (VEGF) signaling. We further exploit our 3D microfluidic system by engineering *in vitro* parabiosis-on-a-chip model through modulation of cross-circulation of blood and niche factors between young and old VMoCs to simulate both heterochronic blood transfusion and parabiosis. Finally, we validate the response of pro- and anti-inflammatory factors found in animal parabiosis by monitoring cytokine levels in real-time. Collectively, our new approach provides a paradigm-shifting platform to overcomes the inherent challenges of using *in vivo* parabiosis and offer a plug-and-play model for investigating cell- and non-cell autonomous regulations in a variety of physiological and pathophysiological conditions.

## Results

### Engineering 3D muscle stem cell niche-on-a-chip

Anatomically, quiescent muscle satellite cells (MuSCs) are located juxtaposed in between myofibers and basal lamina (**Figure 1A**). In addition, recent evidence suggests that vascular endothelial cells (ECs) and neuromuscular junctions (NMJ) are also located closely in contact with MuSC and influence its myogenic function **(Figure 1A)**. To capture the anatomical features of MuSC niche, we micro-engineered vascularized muscle-on-a-chip (VMoC) with two polydimethylsiloxane (PDMS) layers separated by a thin polycarbonate porous membrane, stacked on top of the glass coverslip for microscope imaging (**Figure S1A**). The top layer of the chip mimics the luminal space of the vascular endothelium with an EC monolayer on the porous membrane designed to allow the molecular transport of systemic and local paracrine factors. The bottom layer consists of three parallel microfluidic channels that are partitioned by a series of micropillars (**Figure S1B**). The center channel of the bottom layer provides a space to reconstruct the 3D muscle microenvironment with gelatin-based extracellular matrix (ECM)-hydrogel, FACS-purified primary MuSCs, and mature myotubes that mimic myofibers. The muscle cells and hydrogel were introduced into the bottom-center channel and confined by surface tension between hydrophobic micropillars (**Figure S1B**). The bottom-side channels were designed to supply nutrients into the cell-laden hydrogel or to add other niche component cells, such as motor neurons and neuromuscular junction. For the ECM compartment, we developed a gelatin-*g*-hydroxyphenylpropionic acid (GH) hydrogel to simulate biophysical and biochemical properties of native muscle ECM within the device and to recapitulate vascular transport between muscle and circulatory system (**Figure S2A**) (38). This gelling system enabled easy manipulation of elastic modulus of the hydrogel by varying horseradish peroxidase (HRP) and H_2_O_2_ concentrations (**Figure S2B**). Among various hydrogel stiffness conditions (150, 700, and 3000 Pa), we selected the GH hydrogel with a stiffness of 700 Pa for the present study because it provided an appropriate ECM structure for optimal MuSC growth and myotube formation (**Figure S2C**). The mechanical stiffness and molecular microenvironment of myofiber are known to be critical factors in MuSC homeostasis and its self-renewal capacity (39, 40). Since post-mitotic myofibers cannot be cultivated *in vitro*, we substituted native myofibers with C2C12-derived mature myotubes to cope with long-term culture (**Figure 1B**). The previous investigation reported that an underlayer of differentiated C2C12-myotubes exhibits an elastic modulus of *in vivo* myofiber stiffness and facilitates muscle myogenesis (41). To recapture the asymmetric anatomy of quiescent MuSC, freshly isolated MuSCs were seeded on top of myotubes underneath the basal lamina representing GH hydrogel (**Figure S3A**). FACS-purified quiescent MuSCs were isolated from transgenic mice expressing green fluorescent protein (GFP) driven by the β-actin regulatory element (42). GFP^+^ muscle satellite cells, defined as CD45^-^, CD11b^-^, CD31^-^,Ter119^-^, Sca1^-^, CD29^+^, and CD184^+^ cells, have been reported to yield over 95% of Pax7^+^/MyoD^-^ MuSCs (**Figure 1C and S3B**) (11, 25, 42-45). The cellular organization in vascularized muscle-on-a-chip (VMoC) displayed a well-aligned myofiber-like structure in the bottom-center channel, and a tightly connected EC monolayer in the top channel (**Figure 1B**). Additionally, we were able to monitor sequential myogenic events of primary MuSCs in real-time (**Figure 1C)**. Following cell proliferation in growth media for 4 days, reduced serum (2% horse serum) media circulated within VMoC induced MuSC activation and the subsequent myogenic differentiation, showing the multi-nucleated MuSC-derived myotube formation (**Figure 1C, S3A, and Video S1**). Collectively, we successfully engineered physiologically relevant *ex vivo* or *in vitro* MuSC vascular niche within a 3D microfluidic device (**Figure 1D**).

**Figure 1.**
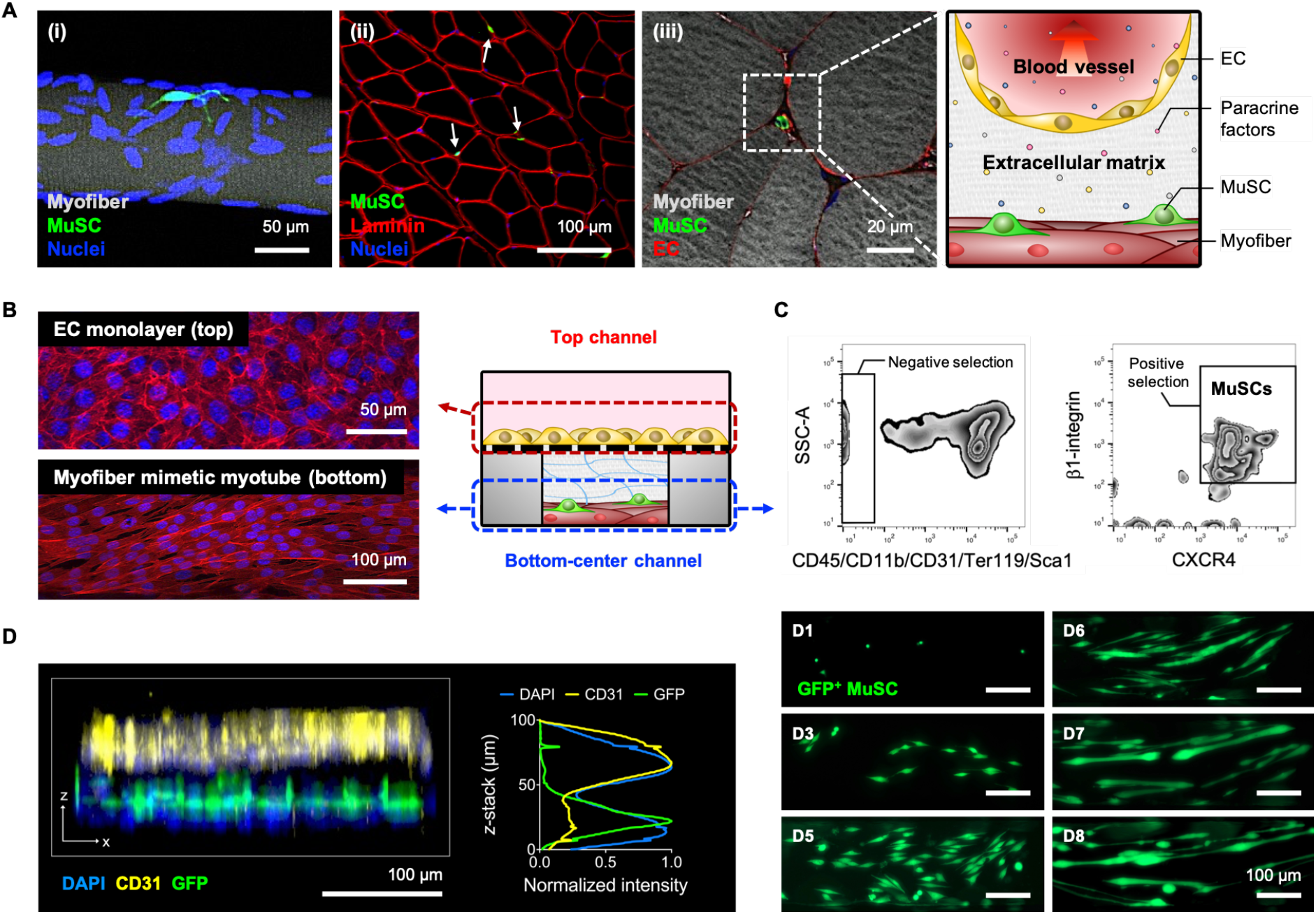
Engineered muscle stem cell niche on a 3D microfluidic platform. (A) Representative images of muscle stem cell (MuSC) microenvironment. Pax7 Cre-R26 flox STOP flox TdTomato mice (MuSC reporter mice; pseudo-colored in green) were used to demarcate quiescent MuSCs. (i) A representative image of Pax7^+^ MuSC on the intact myofiber (actin, gray). (ii) Cross-sectional image of myofibers showing MuSCs and basal lamina (laminin, red). Arrows indicate MuSCs. (iii) Image of MuSC located near capillary endothelial cells (CD31^+^, red). (B) Schematic diagram of vascularized MuSC niche-on-a-chip (VMoC) platform consisting of top and bottom layers compartmented by a porous membrane. Representative DAPI (nuclei, blue) and Phalloidin (F-actin, red)-stained images showing EC monolayer in the top channel and well-aligned myofiber-like myotubes in the bottom-center channel. (C) Representative fluorescence-activated cell sorting plot of MuSCs. Time-course monitoring of FACS-purified quiescent GFP^+^ MuSC proliferation and myotube formation in the VMoC platform. (D) Confocal fluorescence image showing 3D construction of the MuSC niche. The cells were stained with DAPI (nuclei from all types of cells) and CD31 (ECs). The MuSCs express GFP.

### Validation of systemic aging and oxidative stress effects in 3D muscle stem cell niche-on-a-chip

To examine how systemic humoral factors affect survival, proliferation, and myogenic differentiation/fusion of MuSCs within VMoC, we assessed the myogenic activities using the sera isolated from young, old, and *Sod1*^*-/-*^ mice, respectively. Previously, we and others demonstrated that homozygous deletion of an essential antioxidant enzyme, CuZn superoxide dismutase (*Sod1*), accelerated age-dependent muscle atrophy *via* high oxidative damage (46-48). Thus, the *Sod1*^*-/-*^ serum was used to test the systemic oxidative stress on adult myogenesis. The administration of these sera to young MuSCs resulted in distinct myogenic activities (**Figure 2A and 2B**). While young MuSCs cultured in the media containing young serum showed comparable myogenic activities to that in horse serum (HS) treatment (positive control), pro-geronic sera (old and *Sod1*^*-/-*^) significantly reduced myotube formation and the number of fusion events (**Figure 2B)**. In particular, the treatment of *Sod1*^*-/-*^ serum led to a substantial decrease in the relative total cell number, myotube number, and fusion index, suggesting systemic oxidative stress influences myogenic potential.

**Figure 2.**
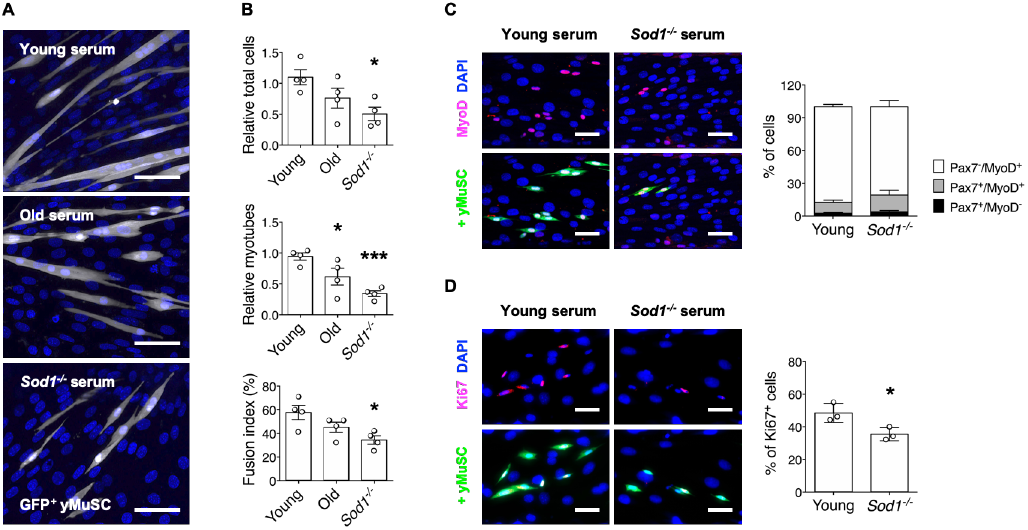
Systemic aging and oxidative stress alter myogenic activities in muscle stem cell niche-on-a-chip. (A) Representative images of DAPI (blue)-stained GFP^+^ young MuSCs (white) grown on top of myofiber-like mature myotubes in VMoC on day 8. Each serum was collected from young (3-4 months), old (20-22 months), and *Sod1*^*-/-*^ (4-6 months) mice, respectively. (B) Quantifications of total cells, myotubes, and fusion index for the young MuSCs (n=4). Relative total cells and myotubes were normalized to each average value obtained from HS treatment (positive control). *\*p<0.05*, and *\*\*\*p<*0.005 *vs*. Young. (C) Representative images of Pax7/MyoD/DAPI-stained GFP^+^ young MuSCs (left) and distribution of Pax7^+^/MyoD^-^, Pax7^+^/MyoD^+^, and Pax7^-^/MyoD^+^ cells cultured in media containing young or *Sod1*^*-/-*^ serum for 2 days (right) (n=3). (D) Representative images of Ki67/DAPI-stained GFP^+^ young MuSCs (left) and quantification of Ki67^+^ cells by young or *Sod1*^*-/-*^ serum treatment (right) (n=3). *\*p<0.05 vs.* Young. All scale bars, 50 μm. All quantitative data are shown as mean ± SEM.

The adult myogenesis is initiated by regenerative cues that activate quiescent MuSCs to enter the cell cycle and express a set of transcriptional factors to commit to myogenic differentiation and to propagate cellular proliferation (6, 49, 50). It has been well documented that aging is associated with oxidative damage to macromolecules (13, 46, 47). To further assess the impact of systemic oxidative stress (*Sod1*^*-/-*^ serum) on MuSC myogenic activation and proliferation, we next compared MuSC behaviors in response to chronic oxidative stress. When MuSCs were treated with young wild-type or *Sod1*^*-/-*^ sera, no significant difference was observed in Pax7/MyoD expression ratio, and the majority of cells (> 81%) were MyoD positive (Pax7^-^/MyoD^+^), suggesting MuSCs were readily committing to fusion-competent myoblast in both conditions (**Figure 2C**). However, when *Sod1*^*-/-*^ serum was administered, a significant decline in proliferation was observed, as measured by a decrease in Ki67^+^ proliferating cells (48.3%), as compared to that in the young wild-type serum treatment (35.3%) (**Figure 2D**). Taken together, using adult myogenesis as an end-point measurement, we validated that the systemic conditions can be accurately controlled with high sensitivity at the microscale, and reliably monitored in our VMoC platform.

### Crosstalk between MuSC and EC via VEGF signaling in VMoC

In the next set of experiments, we examined whether systemic soluble factors can be reliably sampled within our VMoC. Following serum administration, microfluidic samples (8 µL of the conditioned media collected from the luminal region) from the VMoC circulatory system were collected and analyzed. The molecular diffusion across the EC monolayer on the membrane between the bottom and top channels takes at least 4 hours to reach equilibrium (**Figure S4**). Thus, we set the sampling time to 8 hours in each chip incubation, allowing sufficient time for soluble factors to secrete and interact with cells in the VMoC. We compared serum (2%) only controls to young (3-4 months) and aged (20-25 months) with young and old VMoCs, which are composed of MuSCs isolated from young and aged animals with young and aged mice sera administered within the microfluidic system, respectively. ECs, ECM hydrogel, and base myotubes remained constant in all groups. Then, ten cytokines known to affect myogenesis and differentially regulated in animal parabiosis were selected and analyzed using the Luminex multiplex assay. In both young and old VMoCs, overall levels of muscle cytokines showed a higher sensitivity compared to the direct measurement from young and old serum control, suggesting a significant number of soluble factors are released from vascularized muscle stem cell niche construct (**Figure 3A and S5**). Intriguingly, of the ten cytokines analyzed, VEGF levels were undetectable in the serum controls but significantly elevated in both young and aged VMoCs. The VEGF signaling associated with ECs has been highlighted as a key molecular pathway regulating MuSC self-renewal and maintenance (51, 52). Thus, we assessed whether VEGF in VMoC alters the myogenic process. We first confirmed that the VEGF was secreted from both ECs and C2C12-derived myotubes in VMoC (**Figure 3B**). The EC continued to produce VEGF with a gradual increase during the monoculture, whereas the myotubes alone secreted VEGF at a significantly lower level. Interestingly, the co-culture of ECs and myotubes led to a synergistic increase in VEGF production as compared to the sum of each cell monoculture, underscoring the importance of crosstalk between ECs and myotubes during adult myogenesis (**Figure 3B**). To further examine the paracrine effects of the vascular endothelium on myogenic differentiation, we analyzed the VMoCs with or without ECs in the top channel. The co-culture with ECs significantly promoted the myogenic differentiation of MuSCs, which is demonstrated by a higher number of myotubes, fusion index, and increased number of myonuclei in multi-nucleated myotubes, as compared to no EC control (**Figure 3C and 3D**). Furthermore, these enhanced myogenic activities were comparable to recombinant VEGF (rVEGF) protein treatment. Both EC and rVEGF conditions showed higher myonuclei number in the MuSC-derived myotubes than monoculture control, suggesting an increase in fusion events between myoblasts (**Figure 3E**). Our findings support the notion that VEGF secreted by the vascular niche is a crucial factor in MuSC differentiation, and we further validated that our VMoC can accurately recapitulate paracrine interactions within the MuSC microenvironment.

**Figure 3.**
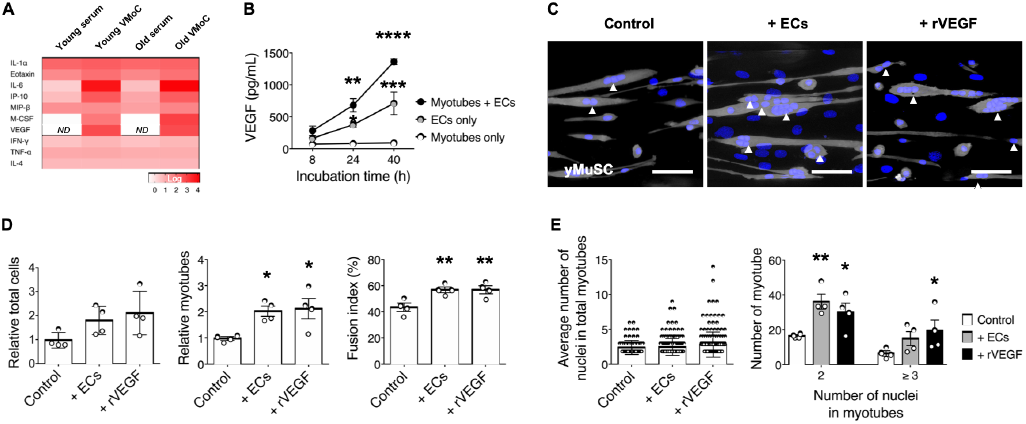
VEGF enhances myogenesis in vascularized muscle stem cell niche-on-a-chip. (A) Mean cytokine levels in the sera from young and old mice (n=9), and the conditioned media from young and old VMoCs (n=3). *ND* (not detected). (B) VEGF levels secreted from myotubes only, ECs only, and co-culture of myotubes and ECs in VMoC system (n=3). \**p*<*0.05*, \*\*\**p*<*0.005*, and \*\*\*\**p*<*0.0001 vs.* Myotubes only at each time-point. (C) Representative images of DAPI (blue)-stained GFP^+^ MuSCs (white) and C2C12-derived myotubes in VMoC on day 8; mono-culture without ECs (control), co-culture with ECs (+ECs), and mono-culture with rVEGF (+rVEGF). Arrowheads (▲) indicate multi-nucleated young MuSC-derived myotubes. Scale bar, 50 μm. (D) Evaluation of young MuSC activities (total cells, myotube number, and fusion index) by co-culturing with ECs or rVEGF in VMoC (n=4). Relative myotubes were normalized to the average value obtained from the monoculture control. \**p*<*0.05*, and \*\**p*<*0.01 vs.* Control. (E) Quantification of average nuclei in total myotubes (left) and distribution of 2 and ≥ 3 nuclei in MuSC-derived myotubes (right). \**p*<*0.05*, and \*\**p* <*0.01 vs.* Control. All quantitative data are shown as mean ± SEM.

### Replicating heterochronic parabiosis and blood transfusion in 3D microfluidic chips

Heterochronic parabiosis, in which young and aged animals are surgically joined to share blood circulation, has been shown to significantly rejuvenate age-acquired defects in muscle function (10, 19, 23, 25). In accordance with prior findings, the age-associated muscle atrophy and fibrosis were attenuated when aged mice were exposed to the young systemic circulation, as confirmed by blood chimerism analysis (**Figure S6**). The heterochronically paired old muscles were rejuvenated, demonstrating an increased diameter of myofibers and reduced fibrosis as compared to the isochronically paired old muscles (**Figure S7**).

Previous studies revealed that both young-blood transfusions to an old systemic environment and heterochronic exchange between young and old blood have shown positive outcomes in rejuvenating cognitive function and restoring the regenerative capacity of aged stem cells (21, 23). However, the blood transfusion does not fully capture the beneficial effects of *in vivo* heterochronic parabiosis. In heterochronic parabiosis, not only blood but also organs, including blood-forming bone marrows, are continuously being shared between young and old animals, constantly replenishing physiological levels of blood exchange and newly formed blood cells, including immune cells. Consequently, one major reason for recent failure in the clinical trial of heterochronic blood transfusion may lie in the lack of our understanding of the underlying mechanisms of frequency, duration, and concentration of blood exchange in heterochronic parabiosis (53). With this in mind, we aimed to engineer an *in vitro* parabiosis model with the capacity to precisely control the exchange of blood-borne factors from the young and old tissues, as well as the blood itself.

To recapitulate both *in vitro* blood exchange and heterochronic parabiosis, we adjusted and synchronized the inter-circulation between young and old muscle microenvironment models (**Figure 4A**). For blood transfusion, the conditioned media from young VMoC were heterochronically exchanged, whereas, in the parabiotic muscle system, the secreted and rejoined factors from young and old MuSC vascular niche were shared. That is to distinguish potential rejuvenation effects of soluble factors in a single-organ muscle system. The parabiotic circulation was designed to test the combinatory effects of young organ sharing on the myogenic function of old MuSCs.

**Figure 4.**
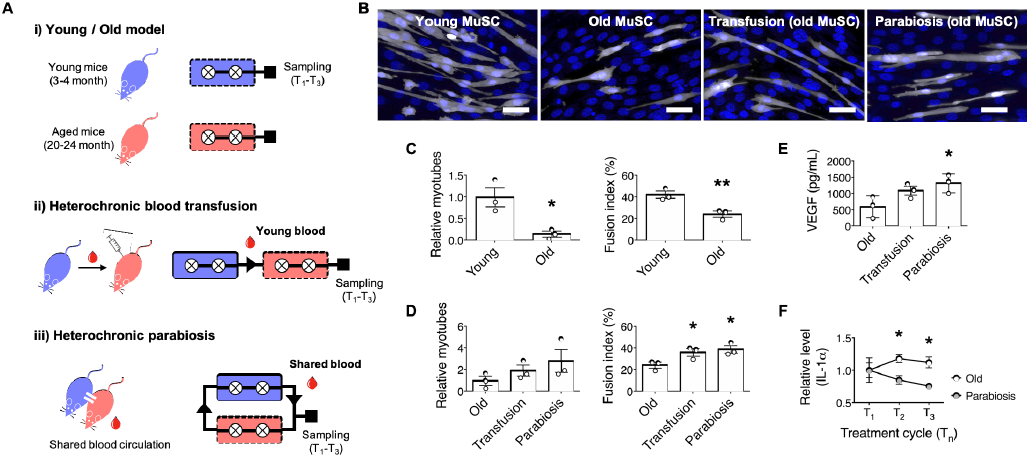
Engineered heterochronic parabiosis-on-a-chip. (A) Diagram of engineered blood-circulation methods for heterochronic blood transfusion and parabiosis. (B) Representative images of DAPI (blue)-stained GFP^+^ young MuSCs/old MuSCs and myotubes after treatment of engineered blood-circulation. Scale bar, 50 μm. (C) Quantifications of myotube number and fusion index for young MuSCs and old MuSCs (n=3). \**p*<*0.05*, and \*\**p*<*0.01 vs.* young MuSC. (D) Old MuSC myogenic activities by young blood-transfusion and heterochronic parabiosis (n=3). \**p*<*0.05 vs.* old MuSC. (E) VEGF levels analyzed from old VMoC microenvironment (Old), young blood-transfused VMoC (Transfusion), and parabiosed-old VMoC (Parabiosis) (n=3). \**p*<*0.05 vs.* old VMoC. (F) The time-course dependent level of IL-1α in old VMoC and parabiosed old VMoC microenvironment (n=3). \**p*<*0.05* vs. old VMoC. All quantitative data are shown as mean ± SEM.

The aged MuSCs exhibited a significant decline in frequency and proliferation (**Figure S8**). In addition, relative myotube number and fusion index of old MuSCs were drastically lower than those of young MuSCs (**Figure 4B and 4C**). These results validate that the age-associated decline in MuSC niche function can be reconstituted in VMoC. When the old MuSCs were treated with the conditioned media collected from young VMoC (young blood transfused old MuSC), significant increases in myotube formation and fusion index were seen (**Figure 4B and 4D**). Similarly, in heterochronic parabiosis circulation, a comparable degree of myogenesis was also observed in old MuSCs (parabiosis old MuSC) (**Figure 4B and 4D**). In this regard, we confirmed that myogenesis-related gene expressions (*Desmin, Myod*, and *Myogenin* normalized to housekeeping gene *β-actin*) were relatively upregulated in old MuSCs treated with the heterochronic parabiosis circulation (**Figure S9**). Furthermore, in the assessment of pro- and anti-geronic cytokine targets, we identified that VEGF levels were differently regulated in heterochronic parabiosis (**Figure 4E**). As compared to old VMoC control, the heterochronic parabiosis mimetic VMoC showed significantly elevated levels of VEGF production, further verifying VEGF as one of the key myokines responsible for the rejuvenating effects seen in heterochronic parabiosis. Among the previously identified senescence-associated secretory phenotypes (SASP) in muscle aging (54), IL-1α showed gradually declined levels over time (T_1_-T_3_ sampling cycles) in heterochronic parabiosis as compared to that in control (**Figure 4F**). Taken together, our *in vitro* parabiosis platform provides control of inter-circulation and real-time monitoring of molecular responses in the localized MuSC niche, which is relatively difficult to conduct in animal studies.

## Discussion

Here, we have demonstrated our PMoC system that mimics organ-specific parabiosis with precise control of extrinsic blood-borne factors and cells in the microengineered 3D construction of MuSC microenvironment. Our 3D microphysiological system allowed for the mechanistic studies with high-precision control and real-time visualization of cell-cell interactions and dynamics between different MuSC niche units. We modulated young/old MuSCs and systemic serum levels but maintaining other niche components constant to verify the MuSC myogenesis in response to conditional extrinsic stimuli without contributions of other individual cells. In addition, microfluidic sampling in MPoC circulatory system facilitated precise quantitative measurement to identify and validate the responses of pro- and anti-geronic factors in a non-invasive manner.

A growing number of studies suggested that the paracrine function of the vascular niche is closely implicated in stem cell function during tissue repair (8, 55). The cellular communication between ECs and myogenic cells was found to trigger angiogenesis and myogenesis coupling during muscle regeneration (56). While important roles of the vascular network in stem cell niche have been raised in muscle regeneration and homeostasis, how these niche components alter MuSC myogenesis during aging remains poorly characterized (52, 57). Substantial researches have investigated *in vitro* muscle contractility, myogenesis, and neuromuscular junction (NMJ) formation; however, *in vitro* models have yet to incorporate microvasculature in order to better mimic *in vivo* muscle physiology (34, 58-60). Accordingly, we sought to engineer 3D MuSC vascular niche that recapitulates biochemical and biophysical features of the native MuSC vascular microenvironment, with which we identified VEGF signaling-mediated cellular interaction between MuSCs and ECs in MuSC myogenic differentiation. In the regulated microenvironment with or without EC monolayer, we demonstrated that VEGF paracrine signaling from ECs is crucial to regulate MuSC function.

The heterochronic parabiosis refers to the cross-circulation of humoral factors and cells in multi-organ systems *via* shared blood circulation. During this process, a variety of biological events, including inflammation and immune responses take place in *in vivo* animal models. These dynamic interactions make it difficult to pinpoint the exact mechanism of the putative youthful factor responsible for muscle rejuvenation (24). Therefore, to reduce confounding valuables and simplify the interpretation of systemic interactions, we focused on a single organ (muscle)-specific parabiosis with modulatory inter-circulation between young and old VMoCs. Despite the evidence of heterochronic blood transfusion providing rejuvenation effect of young blood on aged tissues, the results from heterochronic blood exchange showed different outcomes when compared to heterochronic parabiosis in animal studies (23, 29). Moreover, a recent small clinical trial on the transfusion of young blood failed to recapitulate the beneficial effects of heterochronic parabiosis seen in animal experiments (53). The major distinction between two model systems is that heterochronic parabiosis not only cross-circulate young blood into aged tissue but also share young organs that function as the endocrine system. Thus, we intended to mimic both heterochronic blood exchange and heterochronic parabiosis in our MPoC circulatory system to verify the differences in MuSC myogenic function and local/systemic factors’ responses in the localized muscle microenvironment. We found that both continuous exposure of young systemic environment and heterochronic parabiosis circulation restored the myogenic differentiation of aged MuSCs. Intriguingly, this rejuvenation in heterochronic parabiosis was correlated with increased VEGF amount, indicating that VEGF conditioning at the early stage contributes to the rejuvenation of the aged MuSC myogenic function.

Our approach has shown great potential as *in vitro* parabiosis tissue model for screening or identifying putative pro-regenerative factors that might possess anti-geronic properties against muscle aging. Even though our integrated MPoC system can’t fully replicate *in vivo* parabiosis, we believe that it would serve as an *in vitro* model that can complement the limitation of *in vivo* animal model. Unlike conventional *in vivo* and *in vitro* models, our platform can also offer easy on-demand manipulation for various compelling applications through a ‘mix and match’ approach. For example, we can study on 1) NMJ formation and a mechanism by co-culturing motor neurons in the bottom-side channel, 2) age-associated pathophysiological disease process including sarcopenia by combining different aged organs, and 3) therapeutic efficacy of potential anti-geronic factors. In addition, our chip system will overcome some challenges in *in vivo* parabiosis experiment, including surgical operative procedure and high mortality rate caused by vascular anastomosis and wound infection.

Taken together, our innovative MPoC system would address the critical barriers of *in vivo* parabiosis studies and serve as an experimental model to facilitate the understanding of the dynamic regulation of circulating humoral factors and potential drug discovery. More importantly, our approach will be available for human-parabiosis clinical studies by replacing murine cells with human cells, emulating human parabiosis-on-a-chip that blood transfusion cannot recapitulate.

## Materials and Methods

### Materials

Polydimethylsiloxane (Sylgard 184; PDMS) was purchased from Dow Corning (Midland, MI). Gelatin (>300 Bloom, type A from porcine skin), (3-aminopropyl)triethoxysilane (APTES), 3- (4-hydroxyphenyl)propionic acid (HPA), 1-ethyl-3-(3-(dimethylamino)propyl)-carbodiimide (EDC), N-hydroxysuccinimide (NHS), fibronectin from bovine plasma (FN), peroxidase from horseradish (type VI, 250-330 units/mg solid; HRP), and hydrogen peroxide (30 wt.% in H_2_O; H_2_O_2_) were purchased from Sigma-Aldrich (St. Louis, MO). Photoresist (SU-8) and its developer were purchased from MicroChem Corp (Westborough, MA). Polycarbonate membrane (PC) with a pore size of 8 μm was obtained from Sterlitech Corp (Kent, WA). Dimethylformamide (DMF) was purchased from Junsei (Tokyo, Japan). All reagents and solvents were used without further purification.

### Cell culture

Primary muscle satellite cells (MuSCs) were isolated from mouse hindlimb muscles as previously described (61). Mouse myoblasts (C2C12) were cultured in DMEM containing 10% fetal bovine serum (FBS) and 1% penicillin/streptomycin (PS). Immortalized mouse aortic endothelial cells (ECs) were gift from Dr. Hanjoong Jo (Emory University, Atlanta, GA), and cultured on 0.1% gelatin-coated dishes in DMEM with 10% FBS, 1% endothelial cell growth supplement and 1% PS. All cells were incubated under standard culture condition (37 °C and 5% CO_2_).

### Animals

Mice aged between 3 to 4 months were counted as young and mice aged over 20 months were considered as old. Young and Old *C57Bl/6-*β-*actin*-EGFP mice were mostly used to obtain MuSCs in this study were initially provided by Dr. Amy Wagers (Harvard University, Cambridge, MA) and maintained in the Jang lab mouse colony. B6.129(ICR)-Tg(CAG-ECFP)CK6Nagy/J and B6.Cg-*Gt(ROSA)26Sor*^*tm1(EYFP)Cos*^ mice were purchased from the Jackson Laboratory. *Sod1*^*-/-*^ mice were initially provided by Dr. Holly Van Remmen (Oklahoma Medical Research Foundation). Female mice were used in this study. All mice were housed, aged, and/or bred in the specific pathogen-free condition in Physiological Research Laboratory (PRL) at Georgia Institute of Technology. All procedures were performed in accordance with the approval of the Institutional Animal Care and Use Committee (IACUC) at Georgia Institute of Technology.

### Primary muscle satellite cell (MuSC) isolation

MuSCs were isolated through a cell sorting procedure as previously performed (42, 61). Briefly, hindlimb muscle tissues were harvested from young (2-4 months of age) and aged (20-25 months of age) mice, and then they were incubated in 20 mL of DMEM media containing 0.2 % collagenase type II and 2.5 U/ mL dispase for 90 min at 37 °C. After tissue digestion, the resulting media were mixed with same volume of stop media (20% FBS in F10), filtered using a cell strainer with a pore size of 70 μm, and then centrifuged (300 *g* for 5 min at 4 °C) (Allegra X-30R Centrifuge, Beckman Coulter, USA) to obtain the myofiber-associated cell pellet. The cells pellets were washed with HBSS containing 2% donor bovine serum (DBS), and the cells were incubated with primary antibodies. For MuSC sorting, a cocktail mixture containing the following antibodies was used: (1) allophycocyanin (APC)-conjugated anti-mouse CD11b (1:200), CD31 (1:200), CD45 (1:200), Sca-1 (1:200), and Ter119 (1:200), (2) phycoerthrin (PE)-conjugated anti-mouse CD29 (1:100), and (3) biotinylated anti-mouse CD184 (1:100). After incubation for 30 min at 4 °C, the primary antibodies-treated cells were washed, centrifuged (300 *g* for 5 min at 4 °C), and then treated with a secondary antibody (Streptavidin PE-Cy7) (1:50) for 20 min at 4 °C. Following propidium iodide (PI) treatment and strainer filtration (70 μm), the MuSCs (negative selection: PI, CD11b, CD45, Sca-1, and Ter119; positive selection: CD29 and CD 184) were isolated using fluorescence activated cell sorting (FACS) (BD FACS Aria III, BD Biosciences, USA). The freshly sorted MuSCs were used without sub-culturing.

### Synthesis and characterization of gelatin–g–hydroxyphenylpropionic acid (GH) hydrogels

GH polymer (phenolic content: 150 μm/g of polymer) was synthesized by using a EDC/NHS coupling reaction as previously described (38, 62). Briefly, the HPA solution (6 mmol) activated with EDC (6 mmol) and NHS (9 mmol) for 30 min was applied to the pre-heated gelatin solution (5 g dissolved in 150 mL of deionized water), and then reacted at 40 °C for 24 h. After the reaction, the resulting solution was dialyzed against distilled water, filtered, and lyophilized to obtain the GH polymer. The phenol conjugation and HPA content (degree of substitution; DS) of GH polymer were characterized by ^1^H NMR spectroscopy (AS400, OXFORD instrument, UK) and UV visible spectrophotometer (V-750 UV/vis/NIR, Jasco, Japan).

GH hydrogel was fabricated by mixing an equal volume of two precursor solutions: GH solutions each containing (1) HRP (0.5 µg/mL) and (2) H_2_O_2_ (0.00065 and 0.0011 wt.% for 2 wt.% GH, and 0.002 wt.% for 5 wt.% GH). To characterize mechanical stiffness of GH hydrogel, the gel (15 µL) was formed between top and bottom plate in a rheometer (MCR-302, Anton Paar, Austria), and time-course elastic (G’) and viscous (G’’) moduli were measured at 37 °C for 5 min in oscillation mode (1% strain and 5 rad/s of frequency).

### Microfluidic device fabrication

The microfluidic device was designed using SolidWorks software (Dassault Systems SOLIDWORKS Corp., USA). The widths of the top, bottom-center, and bottom-side channels were 400 μm, 300 μm, and 200 μm, respectively. The height of all channels was 100 μm. A conventional soft lithography process was performed using silicon wafers patterned with both top and bottom micro-channels (63). Briefly, a mixture of PDMS pre-polymer and curing agent (weight ratio of 10:1) was degassed, poured onto the top channel-patterned wafer mold, and then polymerized at 80 °C overnight. To create the bottom channel layer with a thickness of 250 μm, the wafer was spin-coated with the same PDMS mixture, and it was subsequently cured at 80 °C for 1 hour. Finally, both PDMS top and bottom channel layers were applied to a plasma treatment for 2 min under vacuum using a plasma cleaner (PDC-32G, Harrick Plasma, USA), and they were subsequently sandwiched by placing a polycarbonate (PC) membrane between the two layers. The membranes with a pore size of 8 μm were reacted in 5% APTES solution for 30 min immediately before use (64). All channels in the fabricated microfluidic device were sterilized using 70% ethanol and incubated at 80°C for 3 days to restore hydrophobicity of the PDMS surface.

### 3D co-culture of C2C12, MuSCs, and ECs in VMoC unit

Prior to seeding C2C12 cells into the bottom-center channel in the device, the FN solution (50 μg/mL in PBS) was injected into the gel filling port, and incubated for 1 h at 37 °C. The channel was washed with PBS, and the C2C12 were subsequently seeded at a density of 1×10^6^ cells/mL. Following 1 h of cell attachment, non-adherent cells were removed by replacing the culture media, and then cultured in DMEM supplemented with 10% FBS and 1% PS. After culturing for 1-2 days, the freshly isolated MuSCs (2-3×10^5^ cells/30 μL) were seeded onto the pre-cultured C2C12 layer, and then incubated for 6 h in the media containing 20% FBS, 2% horse serum (HS), collagen type I (1 μg/mL), laminin (10 μg/mL), 1% GlutaMAX, 1% PS, and basic fibroblast growth factor (50 ng/mL; bFGF). Then, non-adherent cells were gently removed, and the cells were co-cultured for 4 days in DMEM/F10 media containing 20% HS, 1% GlutaMAX, 1% PS, and bFGF. After cell proliferation period, the hydrogel with a stiffness of 700 Pa was formed in the bottom-center channel, followed by EC seeding into the top channel of the device. To establish EC monolayer over porous membrane, the FN (50 μg/mL in PBS) was coated for 1 hour, and then ECs were seeded at a density of approximately 1×10^7^ cells/mL. After cell attachment for 1 h, the media was refreshed to remove non-adherent cells, and then all cells were incubated for another 4 days in differentiation media (DMEM/F10 containing 2% HS, 1% GlutaMAX, and 1% PS). The same media were also introduced to both bottom-side channels (media port). The media were replaced every 8-12 hours. For VEGF group condition, the MuSCs were cultured without EC monolayer in the differentiation media containing recombinant vascular endothelial growth factor (25 ng/mL; rVEGF). For one-round experiment (n=1), we sorted the MuSCs from three mice, and then seeded the cells at fixed cell density (2-3×10^5^ cells/30 μL) into each group chip. All experiments were repeated four times with same condition (n=4).

For systemic serum experiments, each blood was collected from young (3-4 months; male or female), old (20-22 months; female), or *Sod1*^*-/-*^ (4-6 months; female) mice, and their sera were subsequently isolated using a Microtainer tube (BD Biosciences) as per manufacturer’s instruction. Instead of using HS, these sera were respectively diluted with co-culture DMEM/F10 media to prepare both 20% proliferation and 2% differentiation media. All procedures were performed as described above (co-culture protocol of multiple cells).

For Pax7, MyoD, and Ki67 assay, the MuSCs were seeded on the myotube surface, and then cultured in the proliferation media containing young or *Sod1*^*-/-*^ serum in the device. After incubation for 2-3 days, the cells were fixed and stained for Pax7, MyoD, and Ki67. Pax7/MyoD and Ki67 ratios in the MuSCs were analyzed using antibody-stained images.

### In vivo animal parabiosis study

Parabiosis surgery was operated as previously described (19, 25). Briefly, the left or right side of the pair had 30 mm of skin incision on the flank, and 7 mm wound clips were used to connect the mice skin. Then, adjacent elbow and knee joints between the pair were sutured together. Buprenorphine and saline were injected subcutaneously to reduce pain and to prevent dehydration. After 3 weeks, the blood and muscles were separately collected for the following experiments. For blood chimerism analysis, two drops of blood freshly collected from 1) the parabiotic wild-type mouse paired with GFP^+^ mouse, and 2) the wild-type mouse paired with wild type mouse were added to 900 μL of Ammonium-Chloride-Potassium (ACK) lysis buffer in tube. The tube was kept on ice for 10 min., centrifuged (800 *g*, 5 min., 4°C) to obtain the pellet of white blood cells. Finally, the cell suspension in 300 μL of HBSS containing 2 % DBS was analyzed with a Flow Cytometer for GFP^+^ cells.

### In vitro 3D culture of MuSCs in hydrogels

The GH, HRP and H_2_O_2_ solutions were filtered using a syringe filter (0.2 μm) for sterilization. The GH hydrogels (15 μL) containing MuSCs at a density of 1×10^6^ cells/mL were formed in a 48-well plate, and the MuSCs were cultured in the proliferation and differentiation media for 4 days respectively. The morphology of MuSCs cultured in hydrogels with different mechanical stiffness was observed using a fluorescence microscopy (Axio Observer, Zeiss, Germany).

### Molecular diffusion testing

Firstly, GH hydrogel (700 Pa) was formed in the bottom-center channel of the device, and then the ECs were cultured over porous membrane in the top channel. After the bottom-side and top channels were filled with PBS, all inlet/outlet ports were closed to avoid convection-derived flow inside the device. FITC-dextran solution (M.W.: 40 kDa, 0.5 mg/mL) was injected into one of bottom-side channel. As shown in **Figure S4**, the molecular diffusion rate from bottom channel to whole area in top channel was investigated by measuring relative fluorescence intensity (RFU) of the detection area every 15 min. for 4 hrs. using a fluorescence microscopy (Axio Observer.Z1, Zeiss, Germany), and RFU values were obtained by an ImageJ software (NIH, USA).

### In vivo and in vitro immunofluorescence staining

Cells were fixed in 4% paraformaldehyde solution for 30 min. and subsequently washed three times with PBS. For nuclei and F-actin staining, the cells were incubated in PBS containing Hoechst (1:500) and Alexa Fluor 647 phalloidin (1:40) for 1 hour, and they were washed three times with PBS. For immunofluorescence staining of MuSCs and ECs, the fixed cells were blocked and permeabilized overnight at 4°C in blocking buffer solution (2% BSA, 0.5% goat serum, and 0.5% Triton X-100 in PBS). Following three times of washing with PBS, the cells were incubated with primary antibodies overnight at 4°C, washed three times with PBS, incubated with secondary antibodies (1:100) overnight at 4°C, and washed again five times with PBS. We used the following primary antibodies to stain the cells in this study: CD31 (1:40), Pax7 (1:100), MyoD (1:100), and Ki67 (1:50).

The collected Tibialis anterior (TA) muscles from *in vivo* parabiosis were cut to 10 μm thickness using a cryostat (Thermo Scientific Cryostar NX70) and placed them on slides. After fixation with 4% paraformaldehyde solution for 10 min., the tissues were permeabilized in PBS-T (0.1 % Tween 20 in PBS) for 10 min. and incubated in blocking buffer solution (2% BSA, 0.5% goat serum, and 0.5% Triton X-100 in PBS) for 30 min. After that, another blocking process with 5% goat F(ab) anti-mouse IgG H&L was done to prevent non-specific binding of antibodies. The samples were incubated with primary antibody overnight at 4°C; anti-dystrophin (1:250). Following three washes with PBS, the samples were incubated with secondary antibody (1:250) for 2 hours at room temperature and mounted using anti-fading mounting solution with DAPI.

### Myogenic activity analysis of MuSCs

As shown in **Figure S10**, the MuSC-cultured whole area in bottom-center channel of the device were imaged (21 images per chip; with 20× objective) using a fluorescence microscopy (Axio Observer, Zeiss, Germany), and then the MuSC myogenic activities (*i.e.*, total cell number, myotube number, and fusion index) were analyzed. In this study, the multi-nucleated myotube was defined as a myotube with ≥ 2 nuclei. A degree of myogenic differentiation was expressed as the fusion index (%), which was quantified according to the following equation: (*N*_*m*_ / *N*_*t*_) × 100, where *N*_*m*_ and *N*_*t*_ are the number of MuSC nuclei in the myotube and the total number of MuSC nuclei, respectively. Relative total cells and myotubes in data were normalized to the average of total cell number and myotube number from the control group.

### Cross-circulation of media for transfusion and heterochronic parabiosis

Young MuSCs (3-4 months; yMuSCs) and old MuSCs (20-25 months; old MuSCs) were isolated as described above, and the cells (2.5-3.0×10^5^ cells/30 μL) were immediately seeded on the surface of pre-cultured myotubes. The young and old MuSCs were cultured for 4 days in DMEM/F10 media supplemented with 20% HS, 1% GlutaMAX, 1% PS, and bFGF, and the media were refreshed every 8 hours. After 4 days of cell proliferation, the differentiation media containing 2% young or old serum were introduced. As a control (per one cycle), young VMoC (young MuSCs + young serum) and old VMoC (old MuSCs + old serum) were respectively cultured for 16 hours without media cross-circulation. For blood transfusion (per one cycle), the conditioned media collected from young VMoC incubation for 8 hours was cross circulated into old VMoC and then incubated for another 8 hours. For heterochronic parabiosis (per one cycle), the old serum media was incubated in old VMoC and young VMoC for 8 hours each, and then the conditioned media was transferred to old VMoC again. This treatment cycle was repeated three times over 3 days.

After three cycles of treatments, the myogenic activities (*i.e.*, myotube number and fusion index) of the cultured young and old MuSCs were analyzed. At the same time, the conditioned media were collected from every treatment cycles (T_1_, T_2_ and T_3_), and the levels of following cytokines were analyzed using a Luminex multiplexed immunoassay (Millipore): IL-1α, Eotaxin, IL-6, IP-10, MIP-β, M-CSF, VEGF, IFN-γ, TNF-α, and IL-4.

Myogenic gene expressions (*Desmin, Myod, Ckm*, and *Myogenin*) were quantified by qRT-PCR analysis. Briefly, total RNA from MuSCs co-cultured with C2C12 and ECs in the chip (ibidi) were isolated using the RNeasy Mini kit (Qiagen GmBH), and then cDNA was prepared with T100 Thermal Cycler (Bio-Rad, Hercules, CA, USA). To investigate the effect of different blood circulations on the adult myogenesis, standard qRT-PCR was carried out with a StepOnePlus Real-Time PCR system (Applied Biosystems) using TaqMan Fast Universal PCR Master Mix (Applied Biosystems). The results were quantified by the comparative C_t_ method. C_t_ values for samples were normalized to the expression of the housekeeping gene, *β-actin*.

### Statistical analysis

Quantitative data are expressed as mean ± SEM. Statistical analysis was performed using GraphPad Prism 8, and statistical comparisons of analyzed values were obtained from *t-*test, one-way or two-way analysis of variance (ANOVA). For multiple group comparisons, one-way ANOVA with Tukey’s post hoc tests or Dunnett’s post hoc tests and two-way ANOVA with Sidak’s post hoc tests were performed. All tests resulting in *p*-value (<*0.05*) were considered statistically significant.

## Acknowledgments

This work was supported by the National Institutes of Health under award numbers R56 AG063928 (Y.C.J), R03AG062976 (Y.C.J.), R21AR072287 (Y.C.J.), R21AG056781 (Y.K.), and the National Institutes of Health Director’s New Innovator Award DP2HL142050 (Y.K.). The content is solely the responsibility of the authors and does not necessarily represent the official views of the National Institutes of Health. This work was also supported by the National Research Foundation of Korea (NRF) grant funded by the Korea government (MSIP) (NRF-2018R1A2B2004529). We thank the Physiological Research Laboratory, the core facilities at the Parker H. Petit Institute for Bioengineering and Bioscience, and the Institute for Electronics and Nanotechnology at Georgia Institute of Technology, a member of the National Nanotechnology Coordinated Infrastructure, which is supported by the National Science Foundation (ECCS-1542174).

## Supplementary Information

**Figure S1.**
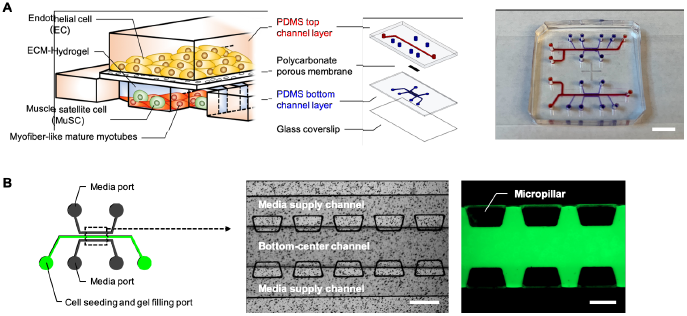
(A) Schematic diagram of vascularized MuSC niche-on-a-chip (VMoC) platform consisting of top and bottom layers compartmented by a porous membrane. The MuSC vascular niche was recapitulated in the device by incorporating ECM-hydrogel and muscle tissue-relevant cells (myofiber-like mature myotubes, FACS-purified quiescent MuSCs, and ECs). A photograph of the VMoC device consisting of two platforms (right). Top channel, red; bottom channel, blue. Scale bar, 5 mm. (B) Design of bottom layer in VMoC. Phase-contrast image of three parallel channels partitioned by a series of micropillar (left). Scale bar, 300 μm. Images after injection of FITC-dextran (green)-loaded hydrogels into the bottom-center channel (right). Scale bar, 200 μm.

**Figure S2.**
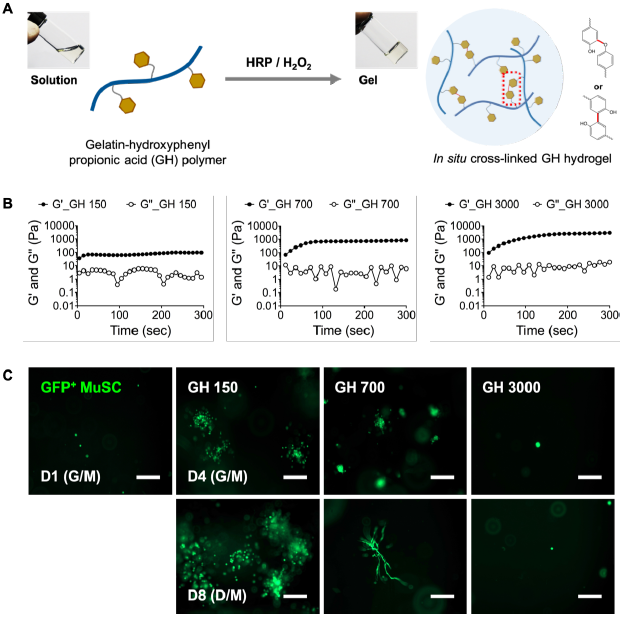
(A) Schematic illustration of *in situ* cross-linked gelatin-*g*-hydroxyphenylpropionic acid (GH) hydrogels formed by HRP/H_2_O_2_-mediated oxidative reaction. The phenol-functionalized GH polymers are catalyzed by the interaction between HRP and H_2_O_2_ and undergo *in situ* cross-linking for hydrogel formation. (B) Time-course measurements of viscous (G’’) and elastic (G’) moduli of GH hydrogels with different H_2_O_2_ concentrations (0.00065 wt.% for GH 150; 0.0011 wt.% for GH 700, and 0.002 wt.% for GH 3000). (C) Representative images of GFP^+^ MuSCs encapsulated in the GH hydrogels. The cells were incubated in growth media for 4 days, then in differentiation media for another 4 days. Scale bar, 200 μm.

**Figure S3.**
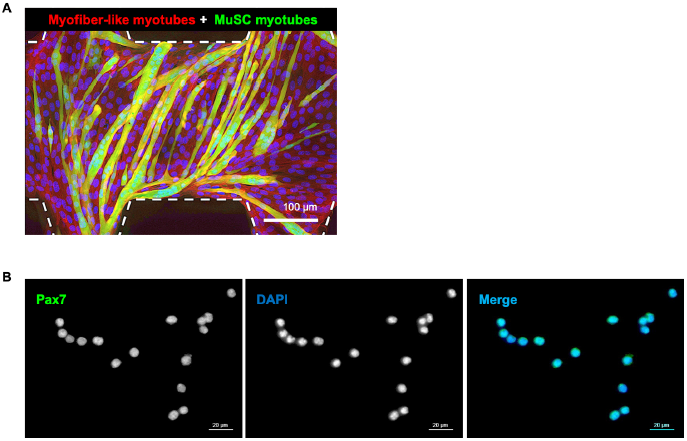
(A) A representative DAPI (nuclei, blue) and Phalloidin (F-actin, red)-stained image showing adult myogenesis of GFP^+^ (green) MuSCs in the bottom-center channel. (B) A representative image of FACS purified quiescent muscle stem cell expressing Pax7 (green) and nuclei (DAPI in blue) at Day 0.

**Figure S4.**
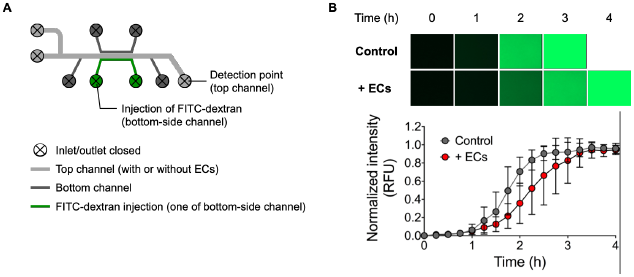
(A) A schematic illustration of experimental set up for diffusion test. The FITC-dextran solution was injected into one of the bottom-side channels, and then fluorescence signal was detected at one of outlet in the top channel. (B) Time-course normalized fluorescence intensity in the condition with or without ECs (n=3).

**Figure S5.**
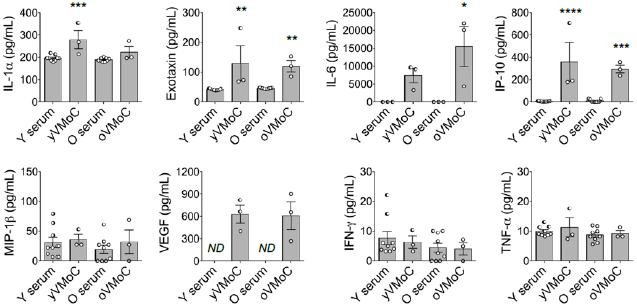
Quantifications of cytokine levels in animal sera and conditioned media from VMoC culture. Young and old sera were collected from young (2-4 months of age) and old (20-25 months of age) mice, respectively (n=9). The conditioned media from young VMoC and old VMoC were collected after 8 h of chip culture (young VMoC and old VMoC) (n=3). \**p*<*0.05*, \*\**p*<*0.01*, \*\*\**p*<*0.005*, \*\*\*\**p*<*0.001 vs.* Young or Old Serum.

**Figure S6.**
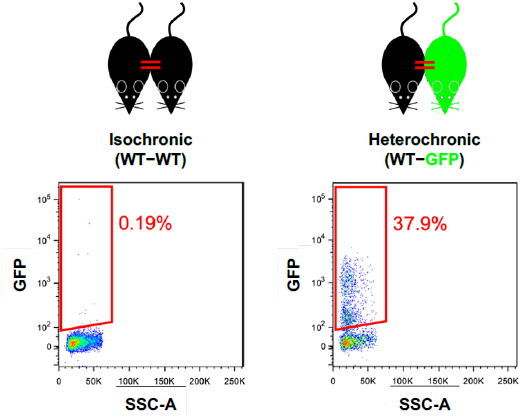
Blood chimerism analysis in a subset of isochronic and heterochronic parabiotic pairs. Flow cytometry of blood cells from the wild-type mouse paired with the wild-type mouse (WT-WT) and the wild-type mouse paired with the GFP^+^ mouse (WT-GFP) after 3 weeks of surgical joining.

**Figure S7.**
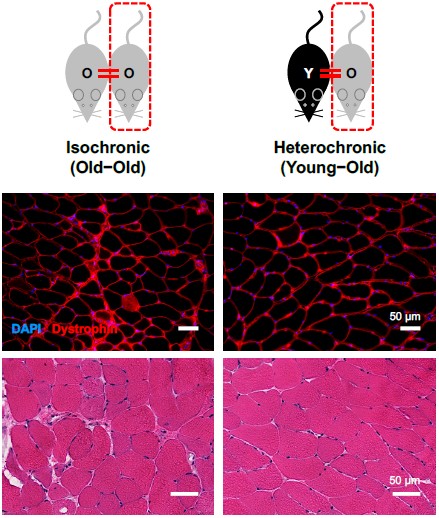
Isochronic aged and heterochronic aged muscle after 3 weeks of parabiosis. (A) Representative images of muscle depicted by DAPI (nuclei) and dystrophin-labeled myofibers. (B) Hematoxylin and eosin (H&E) staining images of isochronically and heterochronically paired old muscles.

**Figure S8.**
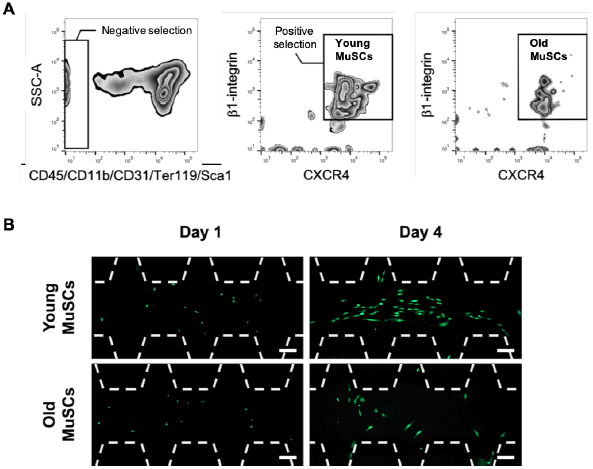
Young and old MuSC sorting plots and proliferation in VMoC. (A) Representative fluorescence-activated cell sorting plot of young and old MuSCs. The FACS-purified quiescent young and old GFP^+^ MuSCs were isolated from young (3-4 months of age) and aged (20-24 months of age) mice, respectively. (B) Representative images show young MuSCs cultured with young serum and old MuSCs cultured with old serum on day 1 and 4. Scale bar, 100 μm.

**Figure S9.**
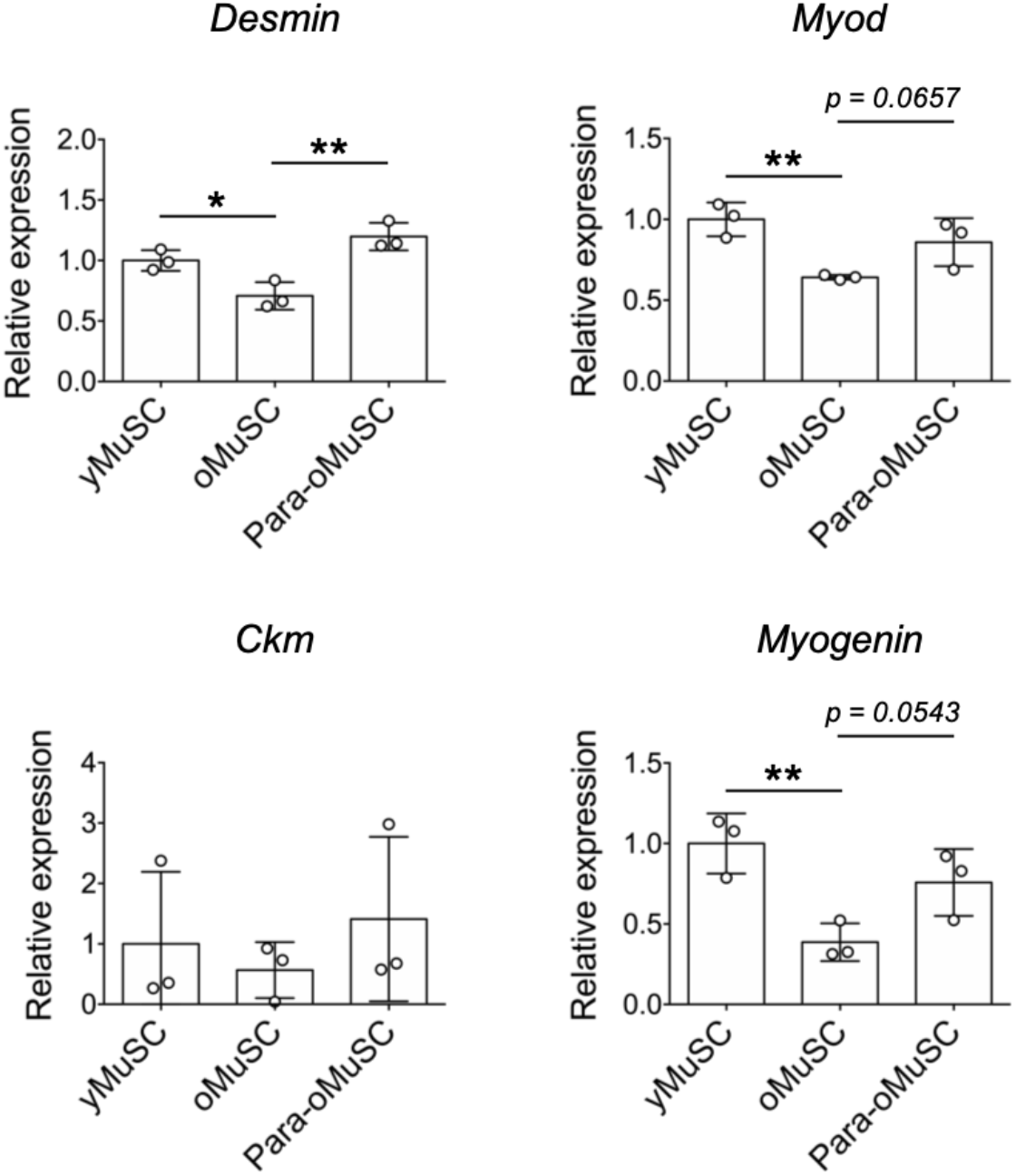
Myogenic gene expressions from MuSCs cultured in young, old, and parabiosis system. Young/old sera and MuSCs were collected from young (3-4 months of age) and old (20-25 months of age) mice. Myogenic gene expressions were measured *via* quantitative RT-PCR in samples from young, old, and Para-old VMoC (n = 3). ^*^*p*<*0.05*, ^**^*p*<*0.01 vs.* young MuSC or old MuSC.

**Figure S10.**
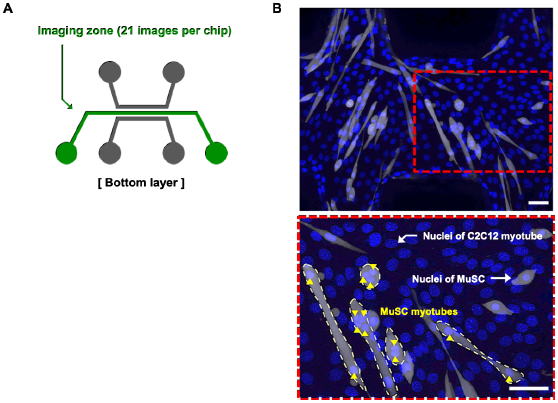
Analysis method of MuSC myogenic activity. (A) Imaged area of MuSCs cultured in the bottom-center channel (green). The whole areas were imaged (21 images per chip with a 200X objective) using a fluorescence microscope. (B) Representative image of DAPI (blue)-stained GFP^+^ MuSCs (gray) and C2C12-derived myotubes. The nuclei of GFP positive MuSCs are distinguished from C2C12 nucleus (bottom). The yellow dotted line indicates MuSC myotube with ≥ 2 nuclei, and arrow (▲) indicates the nuclei in the MuSC myotubes.

